# Distinct Spinal V2a and V0d Microcircuits Distribute Locomotor Control in Larval Zebrafish

**DOI:** 10.1101/559799

**Authors:** Evdokia Menelaou, Sandeep Kishore, David L. McLean

## Abstract

Spinal interneurons coordinate adjustments in the rhythm and pattern of locomotor movements. Two prevailing models predict that interneurons either share or hierarchically distribute control of these key parameters. Here, we have tested each model in the coordination of swimming in larval zebrafish by circumferential excitatory V2a and commissural inhibitory V0d interneurons. We define two types of V2a neuron based on morphology, electrophysiology and connectivity. Type I V2as primarily propagate and amplify rhythmic signals biased to interneurons, while type II V2as primarily segregate and expedite patterning signals biased to motor neurons. Distributed control arises by differences in the likelihood of connections within types and the relative weights of connections between them, but not by a strict anatomical hierarchy. Heterogeneity among V0d neurons supports a similar functional distinction. Our findings provide a hybrid conceptual framework to better understand the origins of rhythm and pattern control in the spinal cord.

## INTRODUCTION

During locomotion animals adjust the frequency, phase and amplitude of rhythmic propulsive movements to navigate effectively (Borgmann and Buschges, 2015; Hultborn and Nielsen, 2007; Pearson, 2000). In vertebrates, phylogenetically-conserved populations of spinal interneurons with distinct molecular and electrophysiological signatures are responsible for coordinating this fundamental process (Gosgnach et al., 2017; Goulding, 2009; Grillner and Jessell, 2009; Kiehn, 2016). After over a century of study (Brown, 1911), two circuit models with different topological predictions have arisen to explain the spinal basis of locomotor control.

In models based on studies of the axial networks of embryonic tadpoles and adult lampreys, spinal interneurons are homogeneous and share responsibility for coordinating the frequency, phase and amplitude of motor output (Grillner, 1981; Roberts et al., 1986). In this single-layer architecture, ‘unit burst generators’ comprised of interconnected excitatory interneurons generate the rhythm and provide ‘first-order’ interneuron and ‘last-order’ motor neuron drive to control the pattern of motor output (Figure 1A, *left*). In contrast, models based on studies of limb networks in neonatal rodents and adult cats use different interneurons for frequency, phase and amplitude control (Gelfand et al., 1988; McCrea and Rybak, 2008). In this multi-layer architecture, first-order interneurons occupy a ‘rhythm-generating’ layer and last-order interneurons occupy a subordinate ‘pattern-forming’ layer (Figure 1A, *right*).

**Figure 1:**
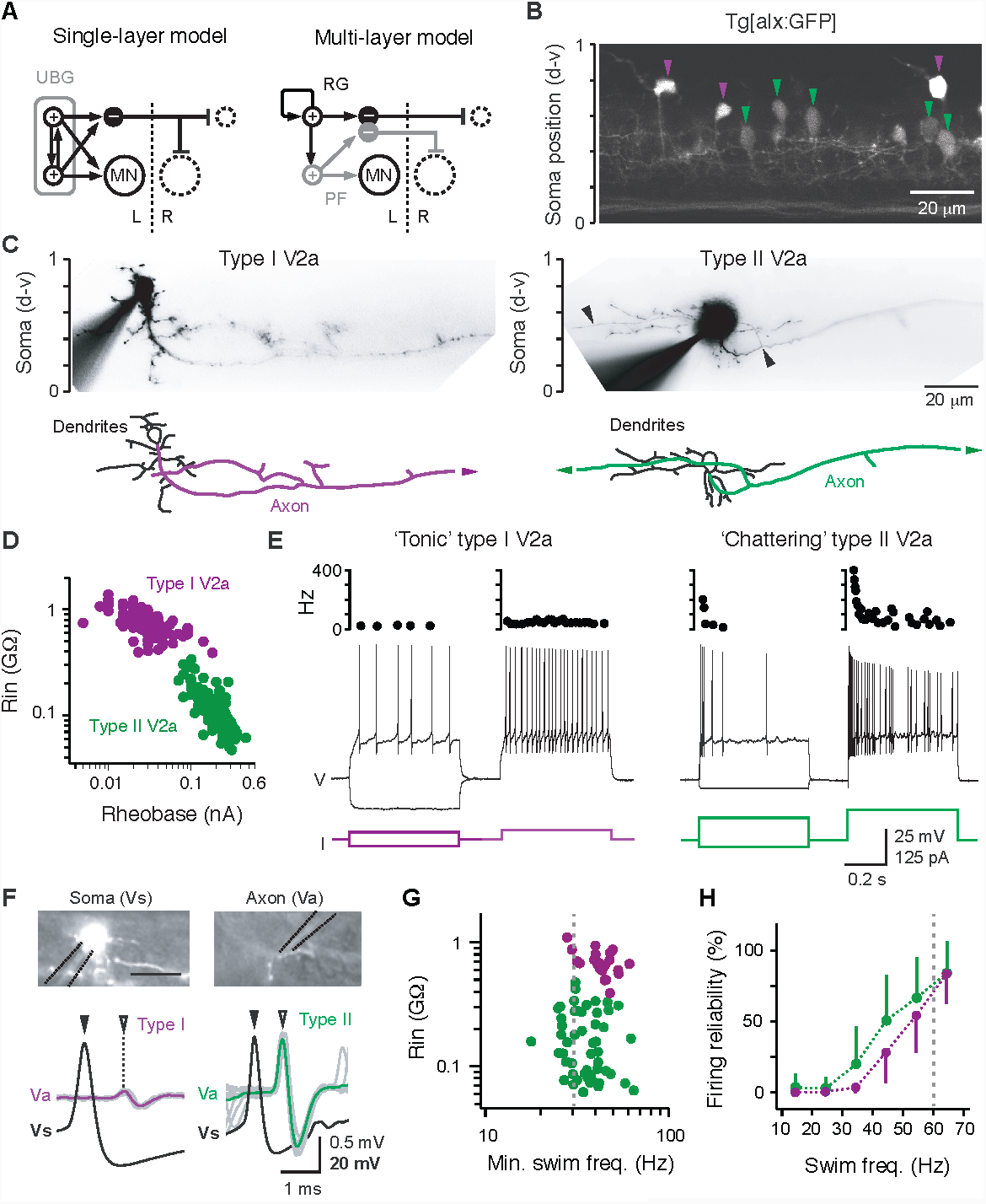
Two types of V2a neurons distinguished by fluorescence intensity, morphology and electrical properties **A**, Schematics highlighting differences in excitatory interneuron (+) connectivity one another, to inhibitory interneurons (–) and to motor neurons (MN) predicted by two prevailing models. UBG, unit burst generator; RG, rhythm generating layer; PF, pattern forming layer. L, left; R, right. Dashed circles indicate contralateral motor neurons (large) and interneurons (small). **B**, Single confocal optical section illustrating type I (purple arrowheads) and type II (green arrowheads) V2a neurons targeted for recordings. Soma positions are normalized to the dorsal and ventral edges of spinal cord (d-v). **C**, Contrast inverted epifluorescent images of fills post-recording (*top*) and reconstructions illustrating differences in the trajectories of the main axon (*bottom*). Type I V2as are descending, while type II V2as are bifurcating. Axon continues out of the field of view (arrowheads). Scale to the left indicates normalized dorsoventral soma position, as in *B*. **D**, Quantification of input resistance (Rin) and rheobase according to V2a neuron type. **E**, *Top*, instantaneous firing frequencies are plotted above the respective traces. *Bottom*, examples of voltage deflections in response to a hyperpolarizing current step (–25 pA for all) and firing responses to current steps around rheobase (*left trace*) and just above (*right trace*) for each V2a type. Resting membrane potentials: type I V2a,–60 mV; type II V2a, –65 mV. V, Voltage; I, current. **F**, *Top*, images of type I V2a neuron stochastically labeled with green fluorescent protein in order to obtain intracellular somatic (Vs) and extracellular axonal (Va) recordings to measure conduction velocities. Dashed lines indicate recording electrodes. *Bottom*, somatic and extracellular axonal recordings from the different V2a types. Somatic spikes are presented as mean waveform (filled arrowheads), while for axonal spikes the mean (open arrowheads) is superimposed on individual traces (grey). See text for quantification of conduction velocity, somatic and axonal spike amplitudes. **G**, Quantification of Rin versus the minimum swimming frequency of recruitment for the different types of V2a neurons. There is no significant relationship despite large differences in Rin (n = 74; Spearman rank correlation analysis, *ρ*(72) = 0.175 p = 0.166). Type I V2a (purple), type II V2a (green). Dashed grey line indicates 30 Hz, below which type II V2a neurons are recruited. **H**, Quantification of the reliability of firing as a function of swimming frequency for the type I (n = 21) and type II (n = 53) V2a neurons. Color code as in *G*. Dashed grey line indicates 60 Hz, when firing in both populations reaches 100% reliability.

Here, we tested the predictions of single-*versus* multi-layer models in the functional organization of spinal locomotor circuitry in larval zebrafish. Studies in zebrafish can bridge work in axial and limb networks, since they swim like tadpoles and lampreys, but provide the genetic accessibility necessary to link molecularly-defined interneuron types to those in mice. In zebrafish and mice different sets of spinal interneurons are used at different speeds of locomotion (McLean and Dougherty, 2015), so we focused on neurons that participate exclusively in high frequency swimming, as they are fewer in number and morphologically identifiable from fish to fish. This also provided an opportunity to definitively assess whether the same interneurons are either first-or last-order or both. We examined two classes of interneuron implicated in spinal locomotor control in zebrafish and mice, namely circumferential excitatory *alx/chx10*-labeled V2a neurons (Al-Mosawie et al., 2007; Ampatzis et al., 2014; Crone et al., 2008; Crone et al., 2009; Eklof-Ljunggren et al., 2014; Kimura et al., 2006; Lundfald et al., 2007; McLean et al., 2008) and commissural inhibitory *dbx1*-labeled V0d neurons (Moran-Rivard et al., 2001; Pierani et al., 2001; Satou et al., 2012; Talpalar et al., 2013), in addition to ‘primary’ motor neurons that innervate either the dorsal epaxial or ventral hypaxial fast-twitch muscle (Menelaou and McLean, 2012; Myers et al., 1986).

We find that V2a neurons can be divided into discrete types. While both types make first-order and last-order connections as predicted by single layer models, complementary differences in their morphologies, electrical properties and patterns of connectivity are consistent with the distribution of rhythm generation and pattern formation predicted by multi-layer models. We also reveal heterogeneity within V0d neurons that supports a similar type of organization. Ultimately, our work reconciles experimental observations in both zebrafish and mice and provides a new conceptual framework to understand the spinal basis of locomotor control and its evolutionary origins.

## RESULTS

### Molecular, morphological and electrophysiological properties define two distinct types of spinal V2a neuron

Our previous work has demonstrated that V2a neurons in zebrafish larvae can be divided into two distinct morphological classes, those with multi-segmental descending axons with extensive local axon collaterals and those with multi-segmental bifurcating axons that can project into the brain (Menelaou et al., 2014). We first assessed how easily we could target these two classes to test the predictions of the single and multi-layer models (Figure 1A). To do so, we performed whole-cell patch clamp recordings of neurons at midbody (segments 10-16) in larval Tg[alx:GFP] fish (Kimura et al., 2006) and targeted neurons occupying more dorsal locations to increase the chances of finding high frequency V2a neurons (McLean et al., 2007; McLean et al., 2008). The transcription factor *alx* is the zebrafish homolog of *chx10*, which labels V2a neurons in mice (Kimura et al., 2006).

Among dorsal V2a neurons, we observed relatively brightly labeled somata displaced from the main V2a population (Figure 1B, *purple arrowheads*) and just below these neurons dimly labeled somata located more laterally (Figure 1B, *green arrowheads*). Post-hoc fills following electrophysiological recordings revealed that brighter GFP-labeled V2a neurons were descending (Figure 1C, *left*), while dimmer GFP-labeled V2a neurons were bifurcating (Figure 1C, *right*). Assuming levels of GFP fluorescence can be used as a proxy for *alx* expression, these observations mirror those in mice, where ‘type I’ V2a neurons with dense local projections exhibit higher *chx10* levels than ‘type II’ V2a neurons, which project both locally and supraspinally (Hayashi et al., 2018). We confirmed differences in GFP intensity were significant by comparing the maximum intensity of bright and dim dorsal V2a neurons in confocal images captured over 4-5 consecutive midbody segments, which revealed an almost two-fold difference in brightness (means, 244 ± 11 a.u. vs. 145 ± 26 a.u., n = 9; Student’s t-test, t(16) = 10.3, p < 0.001). Consequently, to be consistent with mammalian nomenclature we will refer to brighter descending V2as as type I and dimmer bifurcating V2as as type II.

The distinct molecular and morphological features characterizing type I and II V2a neurons were also accompanied by distinct electrophysiological properties. Type I V2a neurons had higher input resistances and lower rheobase values (Figure 1D), consistent with differences in soma size between the types (type I: 25 ± 6 μm2, n = 105; type II: 35 ± 7 μm2, n = 145; Mann-Whitney U-test, U(248) = 1,690, p < 0.001). In response to 500-ms duration direct current steps just above rheobase (1-1.2X), type I V2a neurons generated a stable ‘tonic’ firing response, characterized by spike frequencies ranging from 10–250 Hz (mean, 72 ± 50 Hz, n = 96) with relatively little accommodation (Figure 1E, *left*). Since these frequencies overlap those observed during swimming (McLean et al., 2008), the firing properties of type I V2as could serve as pacemakers or conditional oscillators for rhythmogenesis (Aiken et al., 2003; Buchanan, 1993; Cangiano and Grillner, 2005; Li et al., 2010), which would place them in the rhythm-generating layer.

On the other hand, type II neurons exhibited a more accommodative ‘chattering’ firing pattern, with a barrage of two or more high frequency spikes at the beginning of the step that continued intermittently throughout (Figure 1E, *right*). Instantaneous spike frequencies just above rheobase ranged from 130–650 Hz (mean, 372 ± 99 Hz, n = 140). The higher firing rates and faster adaptation in type II V2as is more appropriate for shaping the duration and amplitude of motor bursts during swimming (Buss et al., 2003; El Manira et al., 1994; Sillar et al., 1992; Sun and Dale, 1998), which is more consistent with occupation of the pattern-forming layer.

Another notable difference between the types was related to their speed of signal propagation. Simultaneous somatic and axonal recordings revealed that somatic spike heights did not differ significantly (Student’s t-test, t(9) = 2.3, p = 0.579), however axonal spikes were significantly smaller (Student’s t-test, t(9) = 2.3, p < 0.05) and conducted more slowly in type I V2as (Student’s t-test, t(9) = 2.3, p < 0.001; Figure 1F). For example, spikes in type II V2as propagated around 0.5 m/s (mean, 0.48 ± 0.04 m/s, n = 4), compared 0.2 m/s (mean, 0.22 ± 0.03 m/s, n = 7) in type I V2as. This is consistent with a larger diameter axon in larger type II V2a neurons (Bawa et al., 1984; Clamann and Henneman, 1976). This also means that once recruited type II V2a neurons would ensure shorter phase lags along the body during higher frequency swimming (Grillner et al., 1976), which is again something more compatible with a pattern-forming function.

### Distinct types of V2a neurons receive distinct patterns of synaptic drive during ‘fictive’ swimming

Given these differences in intrinsic electrophysiological properties, we next examined whether type I and II V2a neurons were recruited at the same swimming frequencies. In response to brief cutaneous electrical stimulation (< 1 ms), short episodes of ‘fictive’ swimming (< 500-ms) comprised of a range of motor burst frequencies are generated (15-80 Hz), with the highest immediately following the stimulus (McLean et al., 2008). Both types of V2a neurons were preferentially engaged at swimming frequencies above 30 Hz (Figure 1G) and firing reliability increased with swimming frequency, reaching 100% above 60 Hz (Figure 1H). Surprisingly, however, larger type II V2a neurons were recruited at frequencies just below 30 Hz (Figure 1G). This is not consistent with the size principle, where smaller neurons should always be recruited at lower frequencies than larger ones. However, it is consistent with reports in mice and older zebrafish, where synaptic inputs to V2a neurons play a more important role in recruitment than membrane properties (Ausborn et al., 2012; Dougherty and Kiehn, 2010).

To see whether differences in their synaptic inputs could explain this disparity, we performed voltage-clamp recordings of excitatory post-synaptic currents (EPSCs) during fictive swimming (Kishore et al., 2014). Both types exhibited rhythmic inward currents corresponding to local ‘fictive’ motor bursts and a slower time course inward current that decreased during the course of the swimming episode (Figure 2A, *left*). On closer inspection, faster time course EPSCs (< 1 ms rise times) were observed superimposed on rhythmic oscillations (Figure 2A, *right*). Among type I V2a neurons, EPSCs were almost exclusively observed at higher swimming frequencies (40-50 Hz; Figure 2B, *top*), while for type II V2a neurons EPSCs were also apparent at lower swimming frequencies (20-30 Hz; Figure 2B, *bottom*). The overall number of EPSCs during fictive swimming at any speed was also greater for type II V2as (Figure 2B), suggesting they receive more excitatory drive. This was also confirmed by measuring peak excitation during fictive swimming, which revealed greater amounts in lower Rin type II V2as (Figure 2C). Consequently, the idea that smaller type I V2a neurons receive weaker, but frequency-specific inputs while type II V2a neurons receive stronger, but more frequency-generic inputs explains their recruitment out of order based on size.

**Figure 2:**
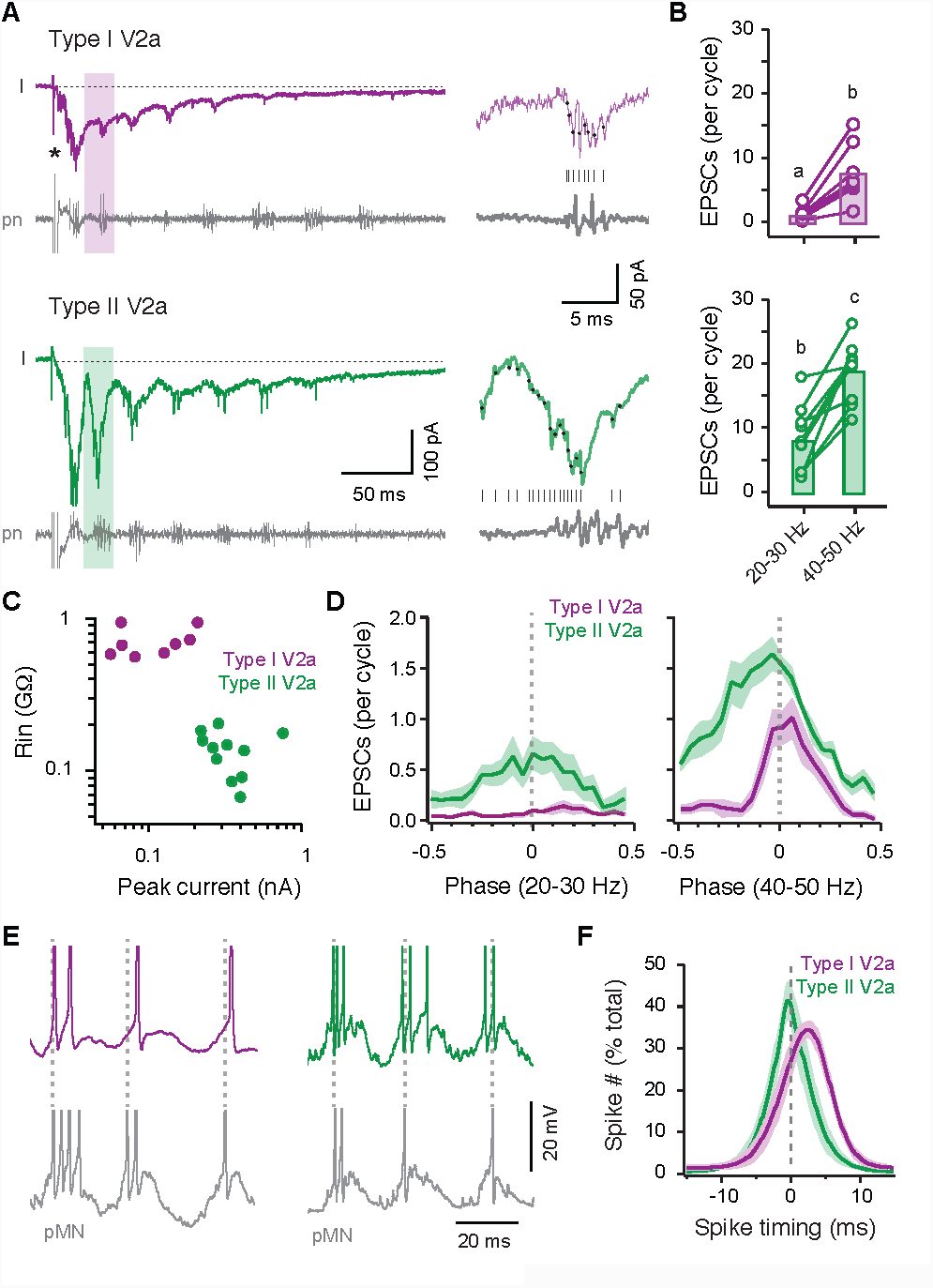
Differences in the levels and patterning of excitatory drive during ‘fictive’ swimming support distinct types of V2a neurons **A**, *Left*, voltage-clamp recordings of excitatory post-synaptic currents (I) and extracellular recordings of peripheral motor nerve activity (pn) measured during ‘fictive’ swimming evoked by a brief electrical stimulus to the tail (artifact at asterisk) from type I (*top*) and type II (*bottom*) V2a neurons. *Right*, individual cycles illustrating EPSC event detection (black dots and lines) relative to proximal motor bursts are taken from regions of the entire swimming episode indicated by shading on the left. **B**, Quantification of the number of EPSCs per cycle at slower (20-30 Hz) and faster (40-50 Hz) swimming frequencies for type I V2as (purple) and type II V2as (green). Means (bars) not sharing letters are significantly different following one-way ANOVA and post-hoc Bonferroni tests (F(3,32) = 26.8, p < 0.001, n = 18). **C**, Quantification of input resistance (Rin) versus the peak excitatory current measured during fictive swimming. Spearman rank correlation analysis, *ρ*(18) = –0.756, p < 0.01, n = 20. **D**, Quantification of the phase of EPSCs with respect to proximal motor bursts (0, dashed line) at slower (left) and faster (right) speeds. Type I V2a, n = 8; type II V2a, n = 8. **E**, Current clamp recordings of excerpts of rhythmic, supra-threshold activity during fictive swimming recorded from type I (left) and type II (right) V2a neurons and a primary motor neuron (pMN) in the same segment. Spikes are truncated to better reveal the underlying synaptic drive. **F**, Quantification of spike number (#) relative to timing of the first spike in a proximal motor neuron (0, dashed line). Gaussian fits of the data reported as mean and standard error. Type I V2a/MN, n = 5; Type II V2a/MN, n = 7.

We next examined the patterning of EPSCs at different frequencies by normalizing the cyclical distribution of EPSCs to phase, where 0 represents the onset of the local motor burst (in-phase). Phase distributions supported the frequency-specific nature of inputs to type I V2as and the frequency-generic nature of inputs to type II V2a neurons (Figure 2D). In addition, while both type I and type II V2a neurons exhibited rhythmic EPSCs that peaked in-phase, in type II V2a neurons EPSCs arrived earlier in the cycle. This suggest that type II V2a neurons are targeted by shorter latency inputs than type I V2as, meaning they could spike earlier and more often, as required for the patterning of motor bursts. To more directly test this idea, we performed paired current-clamp recordings from V2a neurons and primary motor neurons located in the same body segment (Figure 2E). Spike distributions of type I and type II V2a neurons during high-frequency swimming were largely overlapping, preceding and trailing the onset of primary motor neuron activation by ∼10 milliseconds (Figure 2F). Consistent with the voltage clamp analysis, in type II V2a neurons the distribution was skewed toward shorter latency spiking (Figure 2F) and they also fired more spikes per cycle (range, 1-8 spikes; mean, 2.2 ± 1.3, n = 1302 cycles from 54 fish) than type I V2as (range, 1-5 spikes; mean, 1.8 ± 0.9, n = 221 cycles from 21 fish; Student’s t-test, t(1521) = 2, p < 0.001).

The existence of higher impedance V2a neurons with sustained, rhythmogenic firing properties and lower impedance V2a neurons with accommodating firing properties is consistent with work in mice (Dougherty and Kiehn, 2010) and the link between electrophysiological properties to distinct morphological types of V2a neuron is consistent with recent work in older zebrafish (Song et al., 2018). Our data additionally reveal that rhythmic signals carried by type I V2a neurons are frequency-specific and delayed compared to rhythmic signals carried by type II V2as. Collectively, the morphological, electrophysiological and synaptic heterogeneity observed thus far is more consistent with a multi-layer model, where type I V2a neurons occupy the rhythm-generating layer and type II V2a neurons the pattern-forming layer.

### V2a neurons exhibit different patterns of interconnectivity within and between types

Having identified two types of V2a neuron based on levels of GFP expression, morphology, electrophysiology, and patterns of excitatory synaptic drive, we next assessed how connections within and between these types fit the predictions of single-layer and multi-layer models. Given the weight of evidence thus far, our expectation was that rhythm-generating type I V2a neurons would synapse with one another and type II V2as, but not *vice versa*. To assess synaptic connections, we performed paired recordings from identified type I and type II V2a neurons spanning up to four midbody segments. We delivered 5-ms depolarizing pulses that evoked a single spike and analyzed spike triggered individual events and averaged responses.

Within type I V2as, postsynaptic responses were unidirectional, from rostral to caudal, consistent with their descending morphologies (Figure 3A, *left*). The majority of post-synaptic responses were purely electrical (n = 10 of 12; Figure 3D). All paired recordings with electric responses were characterized by inward excitatory post-synaptic currents (EPSCs) recorded at the chloride reversal potential and depolarizing excitatory post-synaptic potentials (EPSPs) recorded at rest with low temporal jitter, relatively fixed amplitudes and no failures (Figure 3C). Additionally, electric responses were sensitive to the gap junction blocker, 18βGA (Li et al., 2009), but not the AMPA-receptor antagonist NBQX (Figure 3E and3F). In the remaining recordings (n = 2 of 12), responses were comprised of mixed electric and glutamatergic components (Figure 3D). Mixed responses were characterized by EPSCs and EPSPs comprised of shorter latency events with low temporal jitter, fixed amplitudes and no failures, and longer latency events with higher temporal jitter and variable amplitudes (Figure 3C). The later component also had relatively high failure rates (65 ± 27%, n = 50). In addition, the earlier events were selectively blocked by 18βGA, while the later events were selectively blocked by NBQX (Figure 3E and 3F). These data are consistent with the idea that type I V2a neurons serve a rhythm generating function, since they exhibit sparse glutamatergic and dense electric interconnections (Kiehn, 2016).

**Figure 3:**
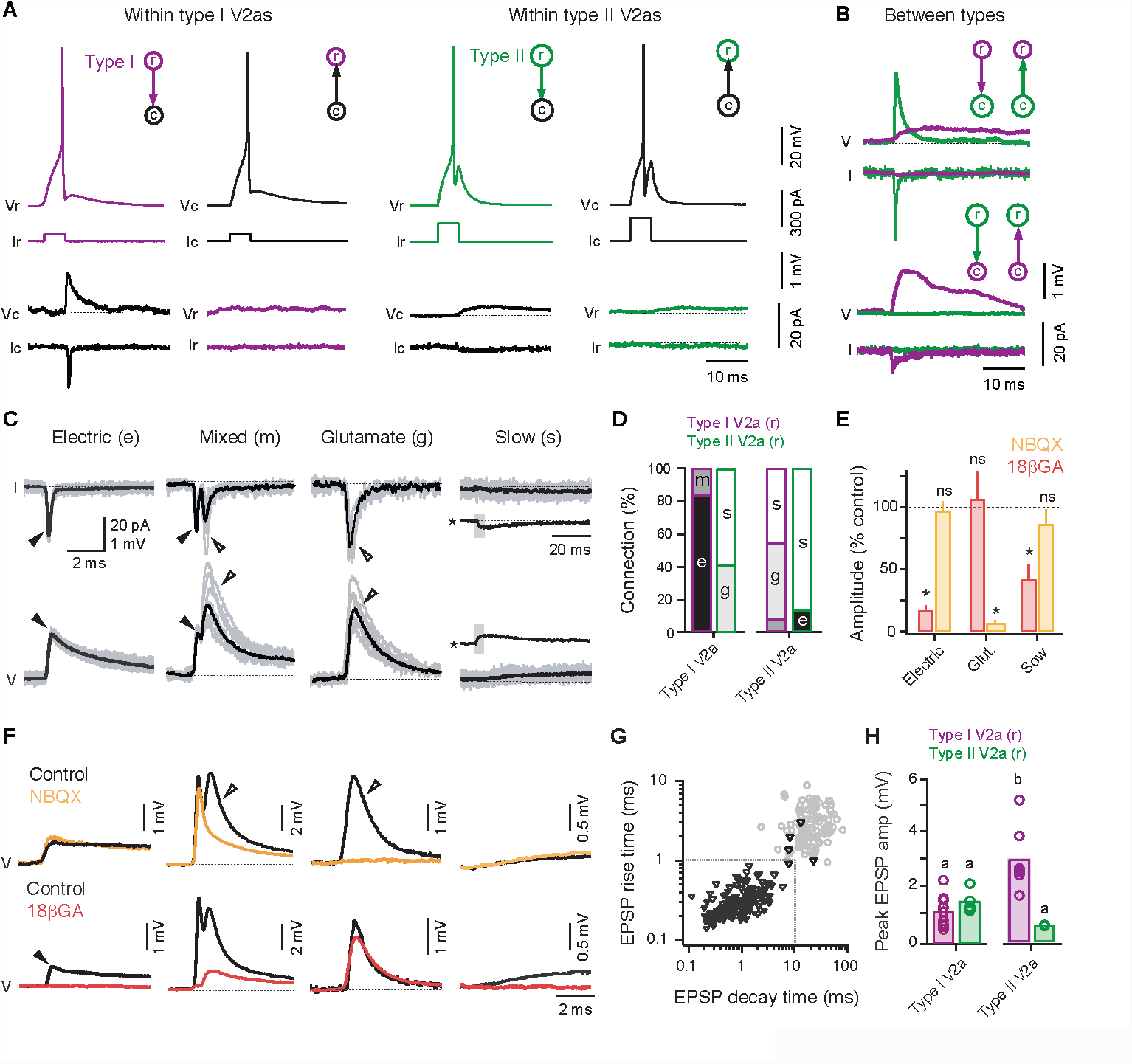
V2a neurons exhibit distinct patterns of connectivity within and between types **A**, *Left*, example of paired recording between type I V2a neurons testing connectivity in the rostral to caudal and caudal to rostral directions. *Right*, example of paired recordings between type II V2a neurons testing connectivity in both directions. Top traces are individual sweeps, while bottom traces are spike triggered averages. I, current; V, voltage. **B**, *Top*, spike triggered averages of post-synaptic responses from a rostral type I V2a neuron to a caudal type II V2a (green) and caudal type II V2a neuron to a rostral type I V2a (purple). *Bottom*, the same for a rostral type II V2a neuron and a caudal type I V2a neuron. **C**, Recordings of spike-triggered post-synaptic events in voltage-clamp (top row) and current-clamp (bottom row). Filled arrowheads mark electrical excitatory post-synaptic currents (EPSCs) and post-synaptic potentials (EPSPs), while open arrowheads mark chemical EPSCs and EPSPs. On the far right, slow electrical EPSCs and EPSPs are also inset on a compressed time scale (asterisks), with a shade grey box indicating the expanded region above or below. Black lines are averages of successful events, grey lines are successful individual events. I, current; V, voltage. **D**, Quantification of the nature of synaptic connectivity within and between the different types of V2a neurons expressed as a percentage of total responses. g, glutamatergic; e, electric; m, mixed; s, slow. Inputs from rostral (r) type I or type II V2as to target caudal neurons on the x-axis are color coded. **E**, Quantification of the effect of drug treatment normalized to control EPSP or EPSC amplitudes for electric, glutamatergic (glut.) and slow responses. Electric 18βGA (*, Student’s t-test, t(16) = 19.9, p < 0.001, n = 9); electric NBQX (ns, Student’s t-test, t(10) = 0.4, p = 0.711, n = 6); glutamatergic 18βGA (ns, Student’s t-test, t(8) = 0.3, p = 0.772, n = 5); glutamatergic NBQX (*, Student’s t-test, t(32) = 38.9, p < 0.001, n = 17); slow 18βGA (*, Student’s t-test, t(8) = 4.6, p < 0.01, n = 5); slow NBQX (ns, Student’s t-test, t(24) = 1.1, p = 0.263, n = 13). Data are reported as mean + standard deviation. **F**, Spike triggered averages of successful post-synaptic responses recorded in current clamp before and after drug treatment. Data are organized from left to right according to type of response, as in *C*. **G**, Quantification of EPSP rise versus decay time for all electrical, glutamatergic and slow responses (n = 191). EPSPs designated as slow are illustrated in grey, while fast EPSPs are in black. Note the 5 outlier fast responses overlapping with slow responses outside the dashed lines are connections from type II V2as to high Rin type I V2as. **H**, Quantification of the peak amplitudes of EPSPs (either electric, mixed or glutamatergic) within and between V2a neurons as in *D*. Means (bars) not sharing letters are significantly different following Kuskal-Wallis ANOVA and post-hoc Mann-Whitney U-tests (H(3) = 17.5, p < 0.0001). Inputs from rostral (r) type I or type II V2as to target caudal neurons on the x-axis are color coded.

Next, we examined functional interactions within type II V2a neurons. Because they have both ascending and descending axons, we expected evidence for reciprocal connectivity. In the majority of recordings (n = 13 of 15), spike-triggered averages revealed bidirectional post-synaptic responses that were relatively slow time course and low amplitude (Figure 3B, *right*). The rise and decay kinetics of these slow responses were an order of magnitude slower than electric and glutamatergic responses (Figure 3G), with rise times as slow as 10 milliseconds and decay times up to 100 milliseconds. While these responses are too slow to explain the cyclical EPSCs observed during fictive swimming, they could contribute to the slower inward current underlying the episode. Slow post-synaptic responses were most sensitive to 18βGA application (Figure 3E and 3F), consistent with an electric connection. In the remaining recordings (n = 2 of 15; Figure 3D), we observed electric responses with faster kinetics, however these were relatively low amplitude (0.19 ± 0.03 mV) and strictly feedforward in nature. These data suggest that type II V2a neurons lack glutamatergic connections and are only weakly electrically interconnected, consistent with a role in pattern formation (Kiehn, 2016).

Surprisingly, however, when we examined the likelihood of connections between types, we found that interactions were not strictly hierarchical, from type I to type II, as predicted by multi-layer models. In recordings when type I V2a neurons were rostral to type II V2as (Figure 3B, *top* and 3D), post-synaptic responses were either glutamatergic (n = 6 of 13), mixed (n = 1 of 13) or slow (n = 6 of 13). All glutamatergic responses, like the glutamatergic component of mixed synapses, were characterized by EPSCs and EPSPs with higher temporal jitter and variable amplitudes (Figure 3C) that were sensitive to NBQX but not 18*β*GA (Figure 3E and 3F). Notably, the failure rate for all purely glutamatergic synapses was significantly higher than the glutamatergic component of mixed synapses (80 ± 18%, n = 47; Mann-Whitney U-test, U(95) = 1,568, p < 0.01). This is consistent with the idea that gap junctions play a role in facilitating chemical neurotransmission at mixed synapses (Liu et al., 2017; Pereda et al., 2004; Song et al., 2016). In contrast, current-evoked spikes in the caudal type II V2a neuron revealed exclusively slow responses in the rostral type I V2a neuron (Figure 3C, *top*).

When type II V2a neurons were the rostral of the two (Figure 3B, *bottom* and 3D), post-synaptic responses in caudal type I V2as were either glutamatergic (n = 5 of 14), slow (n = 7 of 14) or there was no discernable post-synaptic response (n = 2 of 14). Owing to the high Rin of type I V2a neurons, the glutamatergic EPSPs arising from type II V2a neurons had rise and decay times comparable to slow responses (Figure 3G). On the other hand, current-evoked spikes in caudal type I V2a neurons never resulted in discernable responses in rostral type II V2as (Figure 3B, *bottom*). Critically, however, while synaptic interactions between types appear to conflict with a multi-layer model, a comparison of the relative amplitudes of EPSPs demonstrates that type I V2a neurons provide a stronger source of feedforward input to type II V2as than *vice versa* (Figure 3H). When combined with the fact that type II V2a inputs to type I V2a neurons are too slow to pattern their high-frequency activity, these data suggest that type II V2a neurons ultimately rely more on input from type I V2as to shape their rhythmic activity than *vice versa*, consistent with the subordinate role of pattern-forming neurons.

### Indirect electrical continuity is revealed by testing coupling coefficients

The distinctiveness of slow responses and sensitivity to 18βGA prompted us to determine their potential origin. The rise and decay kinetics of slow responses are consistent with low pass filtering of fast events through serial electrical interactions (Luna and Brehm, 2006). Indirect electrical continuity via gap junctions can arise from common target neurons (Eisen and Marder, 1982) and via a shared source of reticulospinal input (Matthews and Wickelgren, 1978). Given that serial electric continuity can complicate assessments of direct synaptic connectivity (Marder et al., 2017), we further explored the nature of electrical interactions within and between types.

To assess non-spike mediated electrical interactions, we delivered 500-ms hyperpolarizing current steps and quantified coupling as a rostro-caudal or caudo-rostral voltage change normalized to the voltage change in the neuron where current was applied (coupling coefficient). As expected based on their dense electric synaptic interconnectivity, we observed bidirectionally symmetric coupling coefficients between type I V2a neurons (Figure 4A and 4E). However, we also found relatively strong bidirectionally symmetric coupling coefficients between type II V2a neurons (Figure 4B and 4E), despite the existence of relatively weak electric connections (cf., Figure 3A, *right*). This is consistent with the possibility that slow responses arise from indirect electrical interactions, which are more easily revealed by more prolonged, hyperpolarizing current steps. To test this directly, we measured coupling coefficients between primary motor neurons separated by four body segments, where there is no chance of a direct physical contact (Figure 4C and 4D). In support, we observed bidirectional coupling coefficients between distal motor neurons of comparable magnitude to that measured for V2a neurons (Figure 4E). These data argue that slow responses are reporting indirect interactions from shared sources of synaptic input.

**Figure 4:**
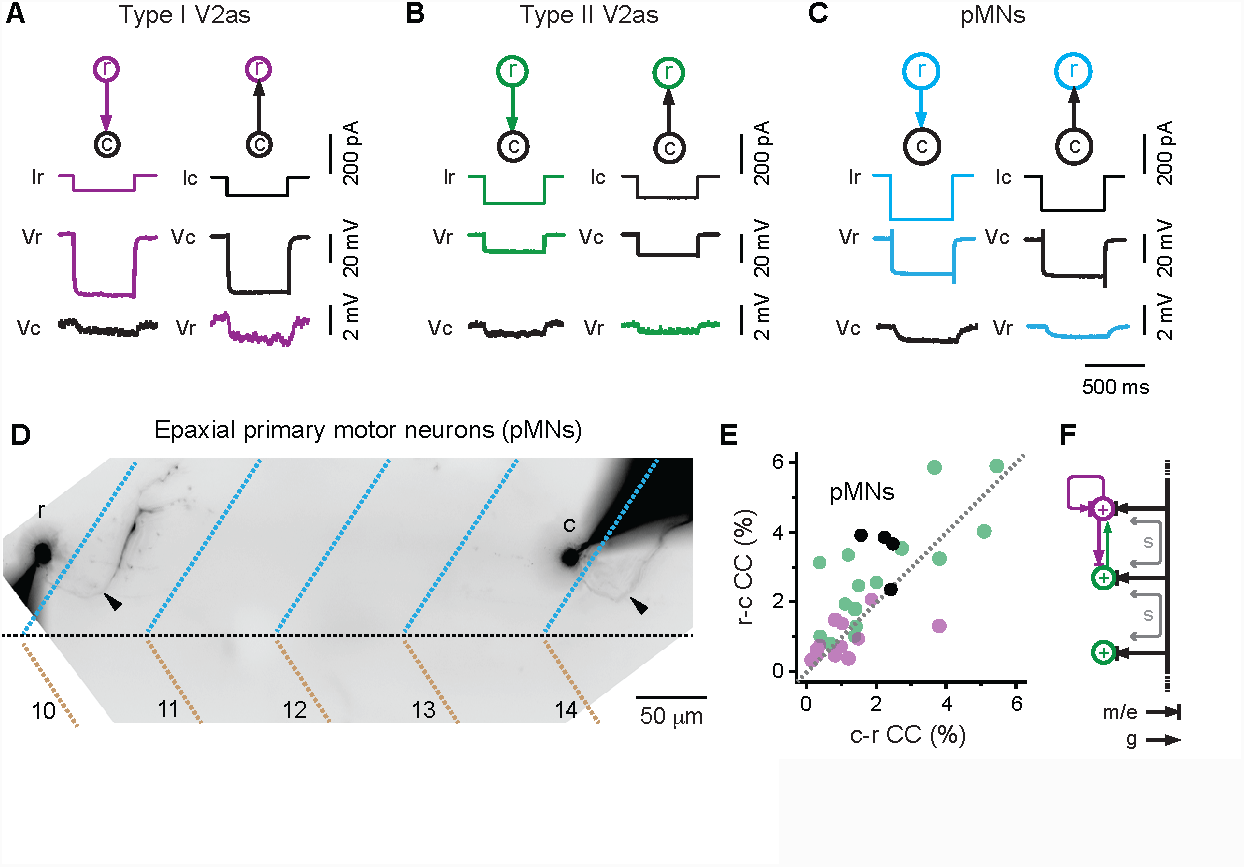
Electrical coupling coefficients reveal indirect electrical continuity **A**, Paired recordings from type I V2a neurons illustrate coupling revealed by hyperpolarizing current steps in both rostral to caudal (r-c) and caudal to rostral (c-r) directions. V, voltage; I, current. **B**, as in A, but for type II V2a neurons. **C**, As in *A*, but for epaxial motor neurons separated by four body segments. **D**, Contrast inverted fluorescent images of fills post-recording of a rostral and caudal epaxial primary motor neurons separated by four body segments, whose coupling is illustrated in *C*. Filled arrowheads indicate axon innervating the dorsal, epaxial muscle (blue dashed lines), which is separated from ventral, hypaxial muscle (brown dashed lines) by the horizonal myoseptum (black dashed line). Body segments are numbered. **E**, Quantification of coupling coefficients (CC) within type I V2as (purple, n = 12 pairs), type II V2as (green, n = 15 pairs) and primary motor neurons (black; n = 2 epaxial/epaxial, n = 1 hypaxial/hypaxial, n = 1 epaxial/hypaxial) in both rostral to caudal and caudal to rostral directions. **F**, Schematic illustrates direct mixed or electric (m/e) and glutamatergic (g) connectivity within and between types, and the indirect nature of electrical continuity observed within type II V2a neurons and between type I and type II V2a neurons that potentially gives rise to slow responses (s). Black line indicates source of common mixed or electrical input.

By focusing on fast synaptic interactions, the picture that emerges is that type I V2a neurons represent a dense electric and sparse glutamatergic circuit that provides stronger, predominantly glutamatergic drive to type II V2a neurons. On the other hand, type II V2as represent a sparse electric circuit that provides weaker, predominantly glutamatergic drive to type I V2a neurons (Figure 4F). This lack of direct interconnectivity and the presence of slow interactions, which most likely reflect indirect electrical continuity (Figure 4F), suggests that individual type II V2a neurons can effectively operate as separate channels. This argues that the distribution of rhythm-generating and pattern-forming functions predicted by multi-layer models is reflected by differences in the likelihood of connections within types and the relative weights of connections between them, but not by a strict anatomical hierarchy.

### Distinct types of V2a neuron exhibit complementary differences in the weight and specificity of last-order connectivity

Previous studies in zebrafish have already demonstrated that V2a neurons are last-order interneurons (Bhatt et al., 2007; Kimura et al., 2006; Song et al., 2016; Song et al., 2018). In addition, we have demonstrated that V2a neurons provide motor pool-specific inputs to allow for separate adjustments in epaxial *versus* hypaxial muscle activity during self-righting responses (Bagnall and McLean, 2014). According to multi-layer models, only the pattern-forming type II V2a neurons should be last-order and provide the drive necessary for pool-specific adjustments in phase/amplitude (Kiehn, 2016). Alternatively, both types could be last-order, with type II V2a neurons having a more dominant and pool-specific influence over motor neurons. To distinguish between these possibilities, we performed paired patch clamp recordings between type I or type II V2a neurons and one of four identified primary epaxial (n = 36) and hypaxial (n = 73) motor neurons.

In contrast to the predictions of multi-layer models, but consistent with a complementary connectivity scheme, both types of V2a neurons formed synaptic connections with primary motor neurons (Figure 5A). Notably, premotor inputs from type I V2a neurons were primarily glutamatergic (n = 31 of 38) or mixed (n = 7 of 38), while inputs from type II V2a neurons were mixed (n = 34 of 71) or electrical (n = 6 of 71). The increased fidelity provided by primarily electric or mixed premotor connections from type II V2a neurons was matched by larger amplitude EPSPs compared to those from type I V2a neurons (Figure 5B). Critically, while we observed evidence for direct premotor connections in all type I V2a recordings (n = 38 of 38), direct connections were only observed in about half of the type II V2a recordings (n = 40 of 71), with the remaining exhibiting indirect slow responses (n = 31 of 71; Figure 5A).

**Figure 5:**
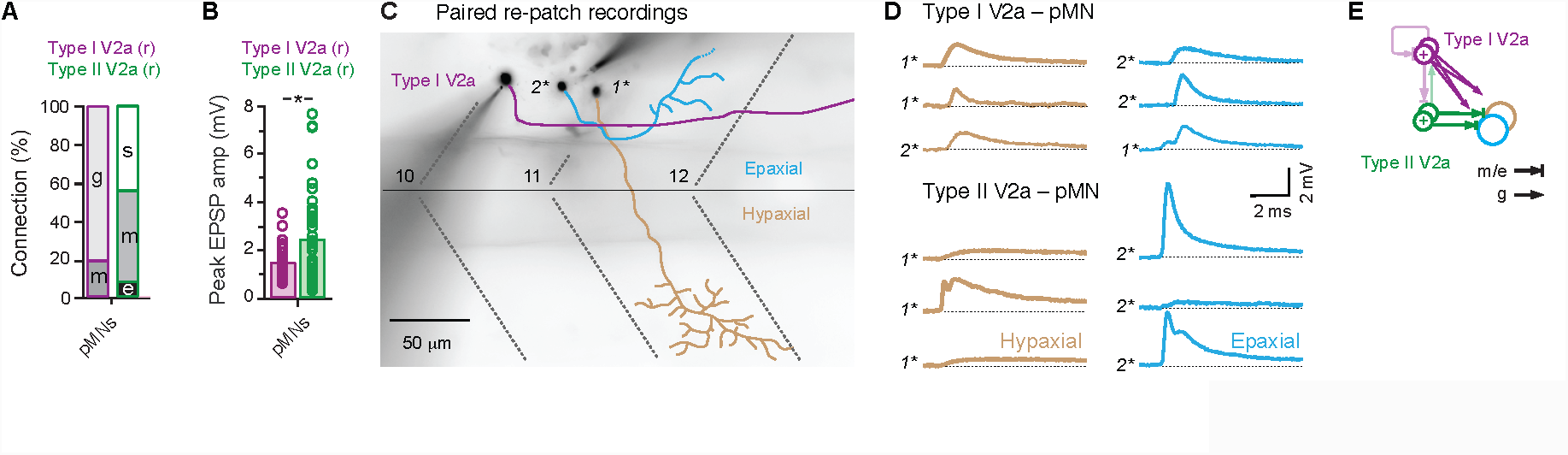
Type II V2a neurons have stronger and pool-specific last-order connections **A**, Quantification of electric, glutamatergic and mixed inputs from type I V2a neurons and type II V2a neurons to primary motor neurons expressed as a percentage of total responses. Inputs from rostral (r) type I or type II V2as to target caudal neurons on the x-axis are color coded. **B**, Quantification of the peak amplitudes of last-order EPSPs (either electric, mixed or glutamatergic) from type I V2a neurons (purple) and type II V2a neurons (green). *, Mann-Whitney U-test, U(76) = 580, p < 0.05). Inputs are color coded as in A. **C**, Contrast inverted fluorescent images of fills post-recording of a type I V2a neuron (purple) and an epaxial (blue) and hypaxial (brown) primary motor neuron. 1*, targeted first; 2*, targeted second. Segment boundaries and the horizontal myoseptum are marked by black dashed lines. Body segments are numbered. **D**, Average excitatory post-synaptic potentials recorded from hypaxial primary motor neurons (brown) and epaxial motor neurons (blue) for type I V2a neurons (top) and type II V2a neurons (bottom). Each row represents a different experiment (n = 3 for each). Order of post-synaptic recording is indicated by 1* (first) and 2* (second). **E**, Schematic summarizes the pool generic glutamatergic (g) last-order connections of type I V2a neurons and the pool-specific mixed/electric (m/e) last-order connections of type II V2a neurons.

The near 0.5 probability of finding direct last-order connections from type II V2as suggests they are the source of the epaxial and hypaxial pool-specific input reported previously (Bagnall and McLean, 2014). To directly test this idea, we performed dual re-patch experiments where epaxial and hypaxial motor neurons were sequentially sampled while holding the same presynaptic V2a neuron (Figure 5C). The majority of these recordings were performed four segments apart (5 of 6 recordings), while only one recording was within the same segment. As expected, for type I V2a to motor neuron recordings when a fast connection was observed in an epaxial motor neuron, a connection was also observed in the hypaxial motor neuron and *vice versa* (Figure 5D, *top*). In contrast, and consistent with pool-specificity, when a connection from a type II V2a neuron was observed in the epaxial motor neuron, only an indirect slow response was observed in the hypaxial motor neuron, and *vice versa* (Figure 5D, *bottom*).

These observations suggest that primary motor neurons receive converging inputs from both type I and type II V2a neurons (Figure 5E). Type I V2a neurons provide a pool-generic frequency signal via lower fidelity glutamatergic synapses, while type II V2a neurons provide pool-specific phase/amplitude signals via higher fidelity electric synapses. In addition to differences in the nature and specificity of connections, individual type II V2a neurons also provide a stronger source of last-order input, which complements the stronger first-order inputs provided by individual type I V2a neurons.

### V0d neurons are also morphologically and electrophysiologically heterogeneous

In multi-layer models, there are inhibitory interneurons that also occupy rhythm-generating or pattern-forming layers commensurate with input from rhythm-generating or pattern-forming excitatory interneurons (Figure 1A). Recent work has shown that glycinergic *dbx1*-positive V0 inhibitory bifurcating interneurons are commissural bifurcating longitudinal (CoBL) interneurons (Satou et al., 2012), which contribute to left-right alternation during swimming in larval zebrafish (Hale et al., 2001; Liao and Fetcho, 2008; McLean et al., 2007). These neurons are homologous to mammalian V0d neurons, which also contribute to left-right alternation during walking (Lanuza et al., 2004; Moran-Rivard et al., 2001; Pierani et al., 2001; Talpalar et al., 2013) and are targeted by V2a neurons (Crone et al., 2008). Consequently, we focused our attention on V0 inhibitory bifurcating neurons, which will refer to as V0d neurons to be consistent with mammalian nomenclature.

We first asked whether heterogeneity exists within V0d neurons to support rhythm-generating and pattern-forming functions. We began by assessing the morphological heterogeneity of V0d neurons, which were transiently labeled by injecting either Dbx:Cre into compound Tg[GlyT2:lRl:Gal4 x UAS:GFP] embryos (n = 14) or a combination of GlyT2:Gal4 and UAS:pTagRFP into Tg[MNET:GFP] embryos (n = 8) and then imaged in 5-6 day old larvae. The latter approach was used to assess putative interactions with primary motor neurons, which are easily distinguished based on their size in Tg[MNET2:GFP] larvae (Balciunas et al., 2004). For purposes of comparison to V2a neurons and primary motor neurons, we focused on relatively ventral V0d interneurons, since these are the ones preferentially recruited during high-frequency swimming (McLean et al., 2007).

Consistent with previous work (Satou et al., 2012), we found three broad morphological types of ventral V0d neurons (Figure 6A): 1) V0d neurons with primarily ascending axons and a descending axon that did not extend beyond two segments (V0d ascending; n = 5); 2) V0d neurons with primarily descending axons and an ascending axon that did not extend beyond two segments (V0d descending; n = 9); 3) V0d neurons with bifurcating axons that each extended beyond two body segments (V0d bifurcating; n = 8). The majority of V0d neurons were unipolar (n = 20 of 22) and had relatively limited dendritic branching compared to V2a neurons. Local axon bifurcations of V0d ‘ascending’, ‘descending’ and ‘bifurcating’ neurons projected dorsally and terminated in close proximity to primary motor neuron somata, while the main axon eventually drifted dorsally to the level of primary motor neuron somata in neighboring segments (Figure 6A, *red ovals*). We also observed occasions where axons projected dorsally, above the motor column or had terminal arbors and en passant connections in between primary motor neurons that could reflect inputs to interneurons (Figure 6A, *open red arrowheads*).

**Figure 6:**
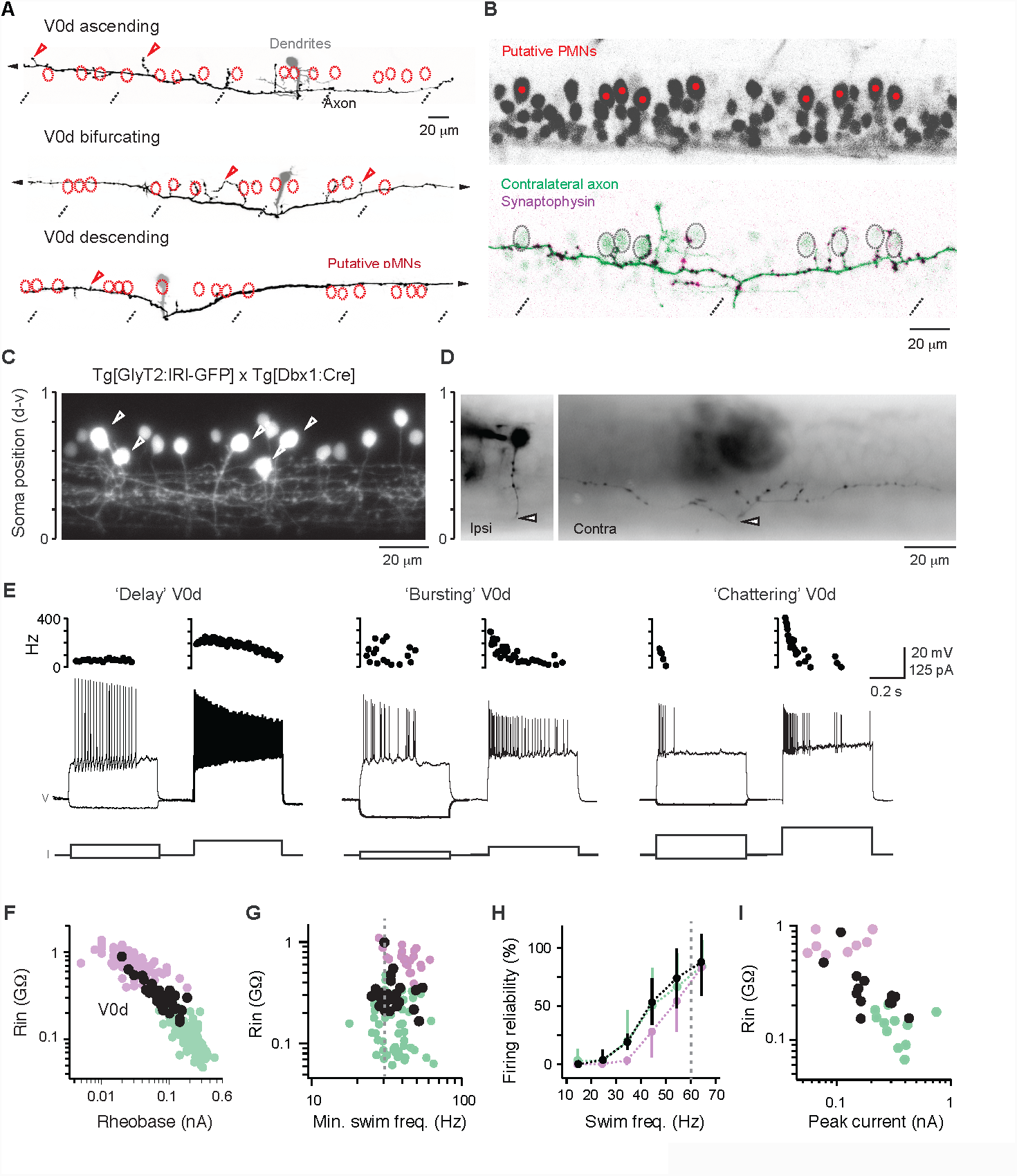
V0d neurons recruited during high-frequency swimming are morphologically and electrophysiologically heterogeneous **A**, Contrast inverted confocal images of ventral V0ds with different morphologies transiently labeled in the Tg[MNET:GFP] line to identify primary motor neurons (pMNs; red ovals). Note fluorescent signals from all motor neurons are not shown for clarity. The longer axon that defines the subtype continues out of the field of view (black arrowheads). Open red arrowheads indicate regions of potential first-order connectivity. The contralateral soma and dendrites are shaded gray. Dashed lines indicate muscle segment boundaries. **B**, *Top*, contrast inverted single confocal optical section of motor neurons in the Tg[MNET:GFP] line. *Bottom*, a single V0d neuron labeled with membrane tagged mCerulean (green) and synaptophysin-pTagRFP (purple). Only the contralateral axon is shown. Dashed ovals indicate the position of putative primary motor neurons (pMNs) labeled by red circles in the image above. Dashed lines indicate muscle segment boundaries. **C**, Single confocal optical section of V0ds targeted for recordings (white arrowheads) in compound transgenic larvae (only GFP fluorescence is illustrated). Scale to the left indicates normalized dorsoventral soma position. **D**, Contrast inverted fluorescent images of a fill post-recording. The contralateral bifurcating axon (origin at white arrowhead) is difficult to track beyond neighboring segments. Scale to the left, as in *C*. **E**, *Top*, Instantaneous firing frequencies are plotted above the respective traces. *Bottom*, examples of voltage deflections in response to a hyperpolarizing current step (–25 pA for all) and firing responses to current steps around rheobase (*left traces*) and just above (*right traces*). Resting membrane potentials: delay V0d, –60 mV; bursting V0d, –63 mV; chattering V0d, –63 mV. V, Voltage; I, current. **F**, Quantification of input resistance (Rin) versus rheobase for V0d neurons (n = 38), superimposed on V2a data from Figure 1D. **G**, Quantification of Rin versus the minimum swimming frequency of recruitment for V0d neurons. There is no significant relationship between the two despite large differences in Rin (Spearman rank correlation analysis, ρ(24) = -0.02, p = 0.895, n = 26). Values are superimposed on V2a data from Figure 1G. Dashed grey line indicates 30 Hz, below which lower Rin V0d neurons are recruited. **H**, Quantification of the reliability of firing as a function of swimming frequency for V0d neurons. Values are superimposed on V2a data from Figure 1H. Dashed grey line indicates 60 Hz, when firing in V0d neurons reaches 100% reliability. **I**, Quantification of Rin versus the peak excitatory current measured during fictive swimming. V0d values are superimposed on V2a data from Figure 2C. Spearman rank correlation analysis, *ρ*(11) = –0.68, p < 0.05, n = 13.

To confirm the synaptic nature of these terminal and *en passant* axonal swellings, we co-injected GlyT2:Gal4, UAS:mCerulean and UAS:Synaptophysin-pTagRFP into Tg[MNET2:GFP] fish. This approach was a relatively low probability one, since it relied on the random incorporation of all three constructs in a single V0d neuron. However, on two occasions we successfully revealed putative synaptic boutons on axon terminals that ramified in close proximity to the somata of primary motor neurons (Figure 6B). Thus, as observed among V2a neurons, morphologically distinct V0d neurons are all likely last-order.

Next, we explored the electrophysiological heterogeneity of V0d neurons. To target V0d interneurons, we used two transgenic lines, either the Tg[GlyT2:GFP] line (McLean et al., 2007) or compound transgenics obtained by crossing the Tg[GlyT2:lRl-GFP] and Tg[Dbx1:Cre] lines (Satou et al., 2012). Again, to simplify our analysis we focused on more ventral and lateral neurons and ignored more dorsal and medial neurons that could participate in lower frequencies of swimming (Figure 6C). Unfortunately, we could not track the commissural bifurcating axon beyond neighboring segments, which precluded links between electrophysiological properties and morphologies (Figure 6D). Nevertheless, we did observe three distinct types of response to direct current injection. ‘Delay’ neurons fired with a delay around rheobase (1-1.2X) and reached instantaneous firing frequencies ranging from 70-350 Hz (mean, 207 ± 94 Hz, n = 7). Firing frequencies throughout the step were stable and demonstrated relatively weak accommodation (Figure 6E, *left*). ‘Bursting’ neurons (Figure 6E, *middle*) fired high frequency spikes around rheobase ranging from 85-600 Hz (mean, 183 ± 177, n = 8) that rode atop slower membrane oscillations at frequencies ranging from 15-40 Hz (mean, 27± 8 Hz, n = 8). We also observed a chattering response similar to the one observed in type II V2a neurons, with higher instantaneous firing frequencies around rheobase ranging from 240-660 Hz (mean, 436 ± 121 Hz, n = 13) that rapidly accommodated (Figure 6E, *right*).

A comparison of rheobase and Rin revealed more continuous clustering of values compared to the discrete clustering of V2a neurons (Figure 6F). There were also no significant differences in either the dorso-ventral distribution (One-way ANOVA, F(2,35) = 0.38, p = 0.682) or sizes of somata (One-way ANOVA, F(2,35) = 0.51, p = 0.604) related to firing type. However, chattering neurons did exhibit significantly higher rheobase (108 ± 30 pA, n = 19; One-way ANOVA, F(2,35) = 5.5, p < 0.01) and lower Rin values (266 ± 60 MΩ, n = 19; One-way ANOVA, F(2,35) = 19.5, p < 0001), than delay (rheobase: 65 ± 20 pA; Rin: 350 ± 118 MΩ, n = 8) and bursting (rheobase: 51 ± 19 pA; Rin: 422 ± 125 MΩ; n = 11) V0d neurons, consistent with observations in V2a neurons.

### V0d neurons can also be distinguished based on distinct patterns of synaptic drive and V2a input

Next, to see if differences in electrophysiological properties of V0d neurons were also matched by differences in excitatory synaptic drive, we performed recordings during fictive swimming. As observed for V2a neurons, lower impedance V0d neurons were recruited at similar minimum swimming frequencies as higher impedance V0ds (∼30Hz; Figure 6G), with 100% recruitment at frequencies above 60 Hz (Figure 6H). Also, as observed for V2a neurons, peak excitatory current was systematically larger in lower Rin V0d neurons (Figure 6I), suggesting the normalization of excitatory drive to neuronal excitability might be a general feature of locomotor circuit organization (Kishore et al., 2014).

We also observed differences in the patterning of synaptic drive during fictive swimming related to electrophysiological properties similar to that observed in V2a neurons. In 8 of 13 V0d recordings, excitation was more apparent at high frequencies (Figure 7A, *top* and 7B), with EPSCs more evenly distributed throughout the cycle, but peaking in-phase (Figure 7C). In the remaining V0d recordings (n = 5 of 13), neurons received comparable levels of excitation at both low and high frequencies (Figure 7A, *bottom* and 7B), with a more narrow distribution of EPSCs peaking in phase with local motor activity (Figure 7C). Critically, V0d neurons with frequency-specific excitatory drive like type I V2a neurons exhibited either delay (n = 4) or burst (n = 4) phenotypes, while the majority of V0d neurons with frequency-generic drive like type II V2a neurons exhibited a chattering phenotype (n = 4 of 5 chattering, n = 1 of 5 burst). The similarity between these observations and those in V2a neurons suggest that V0d neurons are also divided into distinct types for rhythm-generation and pattern-formation. Consequently, for purposes of assessing patterns of V2a connectivity, we divided V0ds into type I delay/burst neurons and type II chattering neurons.

**Figure 7:**
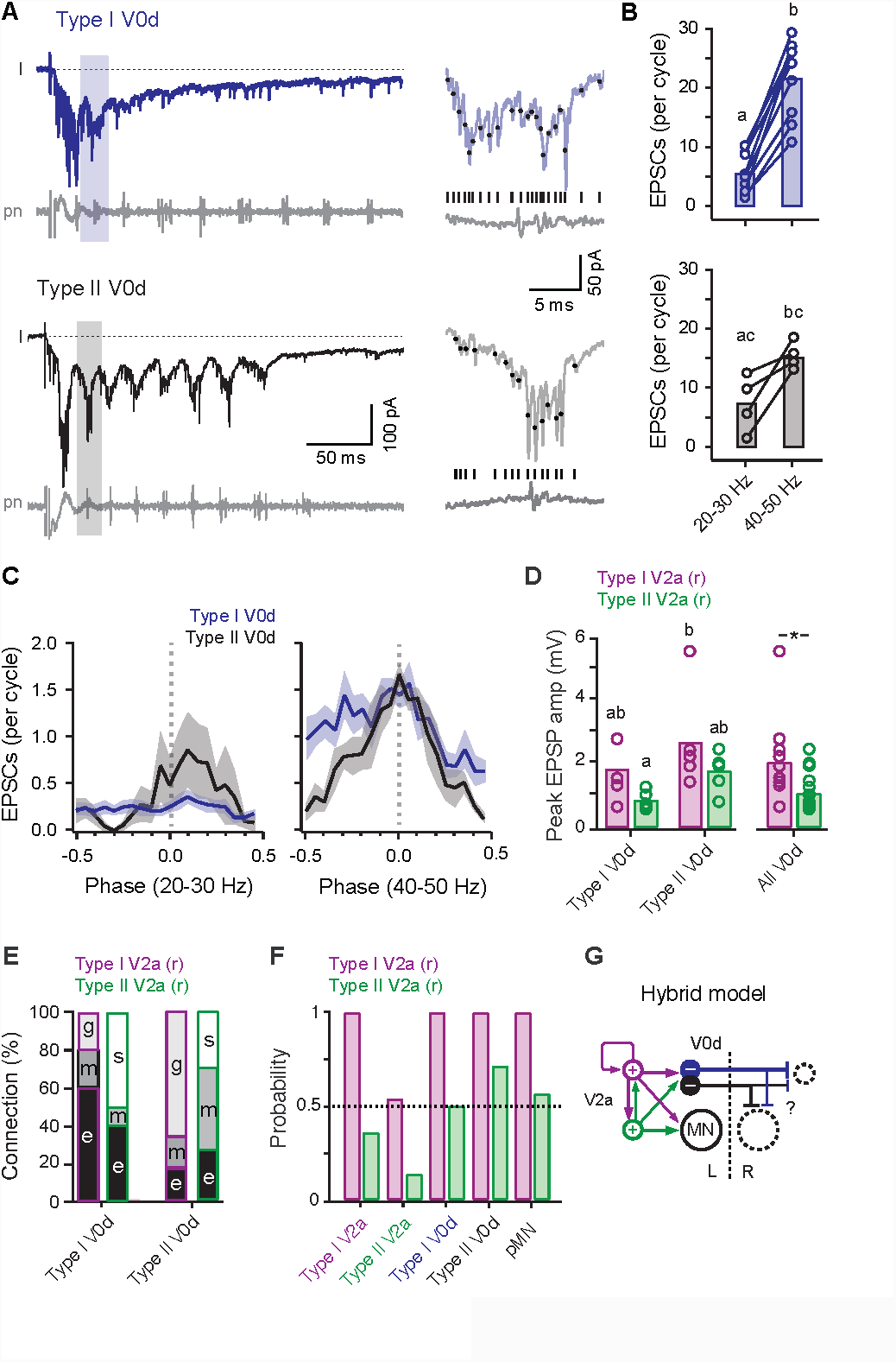
Patterns of synaptic drive during fictive swimming and inputs from V2a neurons support distinct types of V0d neurons **A**, *Left*, voltage-clamp recordings of excitatory post-synaptic currents (I) and extracellular recordings of peripheral motor nerve activity (pn) measured during ‘fictive’ swimming evoked by a brief electrical stimulus to the tail (artifact at asterisk) from type I (*top*) and type II (*bottom*) V0d neurons. *Right*, individual cycles illustrating EPSC event detection (black dots and lines) relative to proximal motor bursts are taken from regions of the entire swimming episode indicated by shading on the left. **B**, Quantification of the number of EPSCs per cycle at slower (20-30 Hz) and faster (40-50 Hz) swimming frequencies for type I V0ds (blue) and type II V0ds (black). Means (bars) not sharing letters are significantly different following one-way ANOVA and post-hoc Bonferroni tests (F(3,32) = 18.8, p < 0.001, n = 13). **C**, Quantification of the phase of EPSCs with respect to proximal motor bursts (0, dashed line) at slower (20-30 Hz; left) and faster (40-50 Hz; right) speeds. **D**, Quantification of the peak amplitudes of V0d EPSPs (either electric, mixed or glutamatergic) from type I V2a neurons (purple) and type II V2a neurons (green). The left two plots separate the data according to V0d type, while the right one reflects pooled V0d data. For separate plots means (bars) not sharing letters are significantly different following one-way ANOVA and post-hoc Bonferroni tests (F(3,17) = 3.2, p < 0.05, n = 21). For the pooled plot*, Student’s t-test, t(19) = 2.1, p < 0.05). Inputs from rostral (r) type I or type II V2as to target caudal neurons on the x-axis are color coded. **E**, Quantification of the nature of synaptic connectivity within and between the different types of V2a and V0d neurons expressed as a percentage of total responses. g, glutamatergic; e, electric; m, mixed; s, slow. Inputs from rostral (r) type I or type II V2as are color coded as in D. **F**, Quantification of the probability of finding a connection from type I V2a neurons (purple) and type II V2a neurons (green) to type I V2a neurons (type I V2a, 12/12; type II V2a, 5/14), type II V2a neurons (type I V2a, 7/13; type II V2a, 2/15), type I V0d neurons (type I V2a, 5/5; type II V2a, 5/10), type II V0d neurons (type I V2a, 6/6; type II V2a, 5/7) and primary motor neurons (type I V2a, 38/38; type II V2a, 40/71). Inputs from rostral (r) type I or type II V2as are color coded as in D. **G**, Schematic of the new, hybrid model highlighting differences V2a interneuron connectivity (+) for control of frequency, phase and amplitude via other V2a neurons, V0d interneurons (–) and motor neurons (MN). L, left; R, right. Dashed circles indicate contralateral motor neurons (large) and interneurons (small). See Figure 1A for comparison to prior models.

Both types of V0d neuron received inputs from type I and type II V2a neurons, with a trend toward stronger inputs from type I V2a neurons, which proved significant following pooled analysis (Figure 7D). There were also differences in the nature and probability of direct connections related to type. Type I V2a inputs were predominantly electric to type I V0d neurons and predominantly glutamatergic to type II V0d neurons (Figure 7E). There was also a probability of 1 in finding a direct connection from type I V2a neurons to both types of V0d neurons, as observed for connections between type I V2a neurons and to primary motor neurons (Figure 7F). Thus type I V2a neurons make dense electric and sparse glutamatergic connections to type I V0d neurons as they make to other type I V2a neurons, and predominantly glutamatergic connections to type II V0d neurons as they make to type II V2a neurons and primary motor neurons. Type II V2a inputs were also predominantly electric to type I V0d neurons, but predominantly mixed to type II V0d neurons (Figure 7E), consistent with their innervation of primary motor neurons.

Notably, type II V2a neurons were more selective in their innervation patterns compared to type I V2a neurons. With the exception of connections to other type II V2as, connections to type I V2a neurons and both types of V0d neurons had a probability around 0.5 (Figure 7F). In the case of motor neuron connection probability, a probability near 0.5 reflects pool-specificity. It is therefore tempting to suggest that pool-specificity among type II V2a neurons is also maintained at the level of interneurons.

Collectively, our findings demonstrate that connections between rhythm-generating V2as and pattern-forming V2as to other V2a, V0d and primary motor neurons are not anatomically hierarchical. Instead, when type I and II V2a neurons converge on common target neurons, first-order interactions are weighted in favor of type I V2a neurons, while last-order interactions are weighted in favor of type II V2a neurons. This complementary scheme contains the heterogeneity and distribution of control predicted by multi-layer models to ensure flexibility in frequency and phase/amplitude control, but with patterns of connectivity predicted by single-layer models. In addition, systematic differences in V2a input related to V0d type and heterogeneity in V0d morphologies, electrophysiological properties and patterns of excitatory synaptic drive are consistent with the distribution of rhythm generation and pattern forming functions within the V0d population. Consequently, the ‘hybrid’ model that emerges provides a new conceptual framework to explain the coordination of rhythm generation and pattern formation during high-frequency swimming in larval zebrafish not only along the body, but also across it.

## DISCUSSION

Our goal was to see how well two prevailing models explained the spinal basis of locomotor control in larval zebrafish. To do so, we took advantage of the identifiable nature and limited number of spinal neurons selectively recruited during high frequency swimming. According to the single-layer model, homogeneous V2a and V0d neurons exhibit first-and last-order connections to control the frequency, phase and amplitude of locomotor output (Figure 1A). For multi-layer models, heterogeneous V2a and V0d neurons are divided into first-order rhythm-generating interneurons that control frequency and last-order pattern-forming interneurons that control phase and amplitude (Figure 1B). Instead of favoring one model, our data support aspects of both models. The resulting hybrid model contains heterogeneous V2a neurons that exhibit complementary patterns of first-and last-order connectivity, with stronger first order connections from type I neurons and stronger last-order connections from type II neurons (Figure 7G). We also propose that V0d neurons are similarly organized based on their morphological, electrophysiological and synaptic heterogeneity (Figure 7G).

Type I V2a neurons fulfill many of the criteria for rhythm-generating neurons. They have dense electric and sparse glutamatergic interconnections, contact participating neurons with high probability, and have intrinsic pacemaking firing properties. Our voltage-clamp recordings from type I V2a neurons at midbody revealed that excitatory drive during fictive swimming is already rhythmic, so pacemaking in these neurons may not be as crucial for rhythm generation *per se*. Assuming more rostral type I V2a neurons exhibit similar intrinsic firing properties (Kimura et al., 2013), they could play a more important role in rhythm generation, as described for escape swimming in tadpoles (Li et al., 2009, 2010). At the very least, type I V2a neurons at midbody play a key role in propagating rhythmic signals. Type I V2a neurons are also high impedance and require little synaptic drive to reach threshold, but they produce large amplitude EPSPs in lower impedance type II V2a neurons and motor neurons. This suggests that type I V2a neurons at midbody also amplify rhythmic signals.

Rhythmic afferent drive to type I V2a neurons during swimming is restricted to high frequencies. This is consistent with the idea that separate sets of V2a neurons carry different frequencies of swimming (Ampatzis et al., 2014). There was also an extremely high likelihood of finding connections from type I V2a neurons to other neurons (100% in most cases). It is unclear if this is due to our focus on neurons recruited at the same frequencies or whether this truly reflects a difference in the density of connections related to speed. Slower circuits are comprised of more numerous V2a and motor neurons (Menelaou et al., 2014) and so may exhibit sparser connectivity. In either case, the relatively small number of dorsal type I V2a neurons involved in high frequency swimming would make them particularly sensitive to experimental perturbations. In support, partial V2a ablation (Eklof-Ljunggren et al., 2012), V2a synaptic silencing (Sternberg et al., 2016), and V2a deafferentation (Knafo et al., 2017) all have the most dramatic effects on high frequency swimming.

Type II V2a neurons fulfill many of the criteria predicted for pattern-forming neurons. They have phasic firing properties better suited for controlling the duration and intensity (i.e., phase and amplitude) of motor bursts. They are also much lower impedance, with faster conduction velocities and a higher likelihood of electric synapses, meaning phase/amplitude adjustments can be transmitted quickly and reliably. Type II V2as are also rarely interconnected and exhibit pool-specific patterns of connectivity, enabling segregated patterning of epaxial and hypaxial motor pool activity (Bagnall and McLean, 2014). While type II V2as rely more on type I V2a neurons to shape their activity than *vice versa*, type II V2a neurons do provide excitatory drive to type I V2as, meaning they can also contribute to rhythm generation. Similarly, type I V2a neurons contribute to pattern formation via type II V2a neurons. While this presumably allows any adjustments in rhythm to be coordinated with pattern and *vice versa*, it also highlights the challenges in relying exclusively on deletion strategies to dissect rhythm *versus* pattern control.

In contrast to type I V2a neurons, type II V2as receive rhythmic excitatory drive at lower swimming frequencies. This suggests they also receive input from slower and/or intermediate type I V2a neurons (Ampatzis et al., 2014; McLean et al., 2008), which would allow for adjustments in phase/amplitude over the full range of swimming frequencies. Adjustments in motor pattern relayed through type II V2a neurons likely originate from fast-conducting reticulospinal, vestibulospinal and propriospinal neurons (Bhatt et al., 2007; Pietri et al., 2009). This would explain why excitatory drive in type II V2a neurons arrives earlier in the swimming cycle than type I V2a neurons and would also explain the earlier arrival of excitatory drive in V0d neurons (Figure 7G). For instance, in cats reticulospinal neurons directly innervate last-order commissural inhibitory interneurons (Bannatyne et al., 2003). The fact that rhythm-generating type I V2a neurons do not always lead local motor activity fits studies of rhythm-generating Hb9 interneurons in mice (Caldeira et al., 2017; Kwan et al., 2009). In this scenario, rhythm-generating neurons are not as critical in initiating locomotor output, instead they provide a gating function, similar to that proposed for mono-synaptic and di-synaptic sensory-motor pathways during limbed locomotion (Buschges and El Manira, 1998).

The morphological, electrophysiological, and frequency-dependent heterogeneity within the V2a population is also mirrored in V0d neurons, supporting their division into rhythm-generating and pattern-forming types. While future experiments are required to map their first-order and last-order connectivity, we would expect complementary differences in the weight and specificity reflecting rhythm-generating *versus* pattern-forming functions as observed in V2a neurons. For example, type II V0d neurons could exhibit stronger, pool-specific last-order connections, while type I V0d neurons could distribute left-right alternating signals to all participating neurons, but with a bias in first-order synaptic strength (Figure 7G). Our anatomical data suggest that all high frequency V0d neurons are at least last-order, which would be consistent with this idea. Also, our previous work has revealed pool-specificity in a proportion of anti-phase inhibitory inputs (Bagnall and McLean, 2014), which could arise from V0d interneurons.

Although past studies have also revealed V2a neurons that use either electric, glutamatergic or mixed synaptic connections (Bhatt et al., 2007; Kimura et al., 2006; Song et al., 2016; Song et al., 2018), our findings additionally suggest that individual V2a neurons of either type can form one or the other based on target neuron identity. For example, type I V2a neurons make primarily electric synapses with other type I neurons, but primarily glutamatergic synapses to type II neurons and motor neurons. Also, type II V2a neurons make primarily mixed synapses with their target neurons, with the exception of type I V2a neurons, which are exclusively glutamatergic. This adds an unappreciated level of complexity to the nature of signal processing by individual spinal excitatory interneurons.

The prevalence of electric synapses is also reported by methods used to calculate electrical coupling coefficients. The coupling we observe here is consistent with type specific coupling among V2a neurons in mice (Zhong et al., 2010) and frequency dependent coupling in older zebrafish (Song et al., 2016). However, what we also demonstrate is that electrical synapses provide an indirect path for injected current to flow. Identified reticulospinal neurons in zebrafish form mixed synapses with motor neurons (Wang and McLean, 2014) and so it is possible that sufficient numbers of these connections provide the necessary electrical continuity. Our previous work also found that electrical coupling is slightly stronger in epaxial and hypaxial motor neurons that share target musculature (Bagnall and McLean, 2014), which could reflect the relative convergence of type II V2a mixed synapses we reveal here. We also report here that type I and II V2a neurons both form electric synapses with V0d neurons, which could provide another indirect source of electrical continuity. In this sense, coupling coefficients can not only reflect direct electric synapses, but also common inputs and shared outputs.

Because we focused on neurons recruited during high frequency swimming, it remains to be seen how a hybrid model explains V2a and V0d circuits controlling lower frequency swimming in larval zebrafish. A recent study in older zebrafish has revealed different classes of V2a neuron based on morphology, firing properties and last-order connectivity (Song et al., 2018). In this case, ‘bursting’ V2a neurons with descending local axons are more likely to make connections to slower motor neurons, while ‘non-bursting’ V2a neurons with local axon bifurcations are more likely to make connections with faster motor neurons. This observation is consistent with the prevalence of intrinsic bursting properties in slower motor neurons (Ampatzis et al., 2013; Menelaou and McLean, 2012) and the idea that V2a neurons with primarily descending axons are also important for rhythmogenesis at low frequencies of swimming. Whether these classes also reflect the type I and type II designations reported here remains to be seen. However, if *alx*-driven GFP levels continue to drop during development as *chx10* levels do in mice (Hayashi et al., 2018), then type II V2as would be challenging to identify in juvenile/adult Tg[alx:GFP] fish.

The ability of our new model to explain locomotor control in larval zebrafish can be summarized as follows. An abrupt stimulus would lead to the activation of low-impedance type II V2a neurons at shorter latencies to provide an initial postural adjustment. If signals are sufficiently robust or persistent they subsequently trigger rhythmic swimming movements via recruitment of high-impedance type I V2a neurons. Type I V2a neurons then provide pool-generic rhythmic excitatory drive upon which is superimposed pool-specific adjustments in phase and amplitude carried by type II V2a neurons. Afferents carrying signals for adjustments in frequency and phase/amplitude could then feed into type I and/or type II V2a circuits, respectively. In this scheme, the ascending axon of type II V2a neurons would serve to relay an internal copy of integrated locomotor commands to brainstem circuitry to gate on-going afferent and/or re-afferent signals (Kozlov et al., 2014).

While the interleaved connections between types would ensure responses to sensory perturbations via type II V2a neurons are appropriately coordinated with propulsive movements, for example when diving or surfacing (Bishop et al., 2016; Ehrlich and Schoppik, 2017; Nair et al., 2015), the relatively sparse connections between type II V2a neurons could also explain how zebrafish generate discrete bilateral dorsal flexions for prey strikes (Hernandez et al., 2002; Patterson et al., 2013), torsional flexions for self-righting responses (Bagnall and McLean, 2014) and unilateral flexions for turning maneuvers (Bhatt et al., 2007; Dunn et al., 2016; Umeda et al., 2016). Thus, a connectivity scheme with parallel type II circuits embedded within a serial type I circuit enables both the segregated and integrated control of rhythmic and discrete movements. Recent work in mice has implicated type II V2a neurons in executing reach and grasp movements (Azim et al., 2014), which would be consistent with their role in discrete movements proposed here. Consequently, the idea that spinal V2a interneurons wired to execute discrete movements can also participate in rhythmic movements is something that could extend to limb control (Alstermark and Isa, 2012; Georgopoulos and Grillner, 1989).

In sum, our work has revealed that distinct types of V2a and V0d neurons effectively distribute control of rhythm-generation and pattern-formation during locomotion in larval zebrafish. During evolution, iterative duplications of this basic circuit motif, one neuron for ‘driving’ and another for ‘steering’, could have supported the coordination of increasingly diverse musculature. In support, similar morphological and synaptic heterogeneity is found among homologous excitatory interneurons in primitive vertebrates, like lampreys (Buchanan et al., 1989; Einum and Buchanan, 2006; Parker, 2003) and the protochordate *Ciona intestinalis* (Ryan et al., 2017; Stolfi and Levine, 2011). Another possibility is that the hybrid framework described here is specific to the evolution of increased aquatic maneuverability in teleosts (Webb and Weihs, 2015), while a true multi-layer framework evolved during the transition to land (Fischer and Witte, 2007), perhaps by eliminating relatively weak connections (i.e., last-order type I V2a synapses and first-order type II V2a synapses). Determining which scenario holds true awaits similarly fine-grained assessments of functional synaptic connectivity in other vertebrates.

## ACKNOWLEDGMENTS

We thank Elissa Szuter for fish care and Eli Cadoff for technical support. We also thank Keith Sillar, Andrew Miri, Joseph Fetcho and members of the lab for feedback. Financial support provided by NIH R01 NS067299 and U19 NS104653.

## AUTHOR CONTRIBUTIONS

Conceptualization, E.M. and D.M.; Methodology, E.M., S.K., and D.M.; Investigation, E.M. and S.K., Formal Analysis, E.M. and S.K., Writing, E.M., S.K., and D.M.; Visualization, E.M. and D.M.; Resources, Supervision and Funding Acquisition, D.M.

## STAR METHODS

### Contactfor Reagent and Resource Sharing

Further information and requests for resources and reagents should be directed to and will be fulfilled by the Lead Contact, David McLean.

### Experimental Model and Subject Details

Adult zebrafish (*Danio rerio*) and their offspring were maintained at 28.5 °C in an in-house facility (Aquatic Habitats). Experiments were performed using 4-5 day old wildtype, Tg[alx:GFP], Tg[alx:lRl-GFP] (Kimura et al., 2006), Tg[MNET2:GFP] (Balciunas et al., 2004), Tg[GlyT2:GFP] (McLean et al., 2007), Tg[Dbx:cre], and Tg[GlyT2:lRl-Gal4;UAS:GFP] zebrafish larvae (Satou et al., 2012). At this stage, zebrafish larvae have fully inflated swim bladders and are free swimming, but have not yet sexually differentiated. All procedures conform to NIH guidelines regarding animal experimentation and were approved by Northwestern University Institutional Animal Care and Use Committee.

### Method Details

#### Plasmid preparation

Transient, mosaic labeling of V2a or V0d neurons was achieved by using the Gal4-UAS system to drive reporter constructs (Koster and Fraser, 2001). DNA solutions were prepared at concentrations between 15 and 25 ng/μL. We generated the UAS:mCerulean and UAS:Synaptophysin-pTagRFP constructs with the Tol2kit (Kwan et al., 2007). For synthesizing UAS:mCerulean, we first amplified mCerulean from CMV-Brainbow (Addgene plasmid 18720) with primers flanked by gateway cloning sites (5’-GGGGACAAGTTTGTACAAAAAAGCAGGCTTACGGAATTAATTCACAGCCACCA, 3’-GGGGACCACTTTGTACAAGAAAGCTGGGTAGAGTCGCGGTGATCTAGAGTC), subcloned into a middle entry vector, and subsequently subcloned to be driven by a 10x element UAS promoter. To generate UAS:Synaptophysin-pTagRFP, we first amplified synaptophysin from UAS:Synatophysin-GCAMP3 (5’-

GGGGACCACTTTGTACAAGAAAGCTGGGTATCAATTAAGTTTGTGCCC, 3’-TCCTTAATCAGCTCTTCGCCCTTAGACACCATCATCTCGTTGGAGAAGGAT) (Nikolaou et al.,

2012)and pTagRFP from UAS:pTagRFP (5’-GAGCCCACATCCTTCTCCAACGAGATGATGGTGTCTAAGGGCGAAGAGCT, 3’-GGGGACCACTTTGTACAAGAAAGCTGGGTATCAATTAAGTTTGTGCCC). The amplified synaptophysin and pTagRFP fragments were then ‘stitched’ together by using overlapping PCR (Hobert, 2002)using primers that were flanked by gateway cloning sites (5’-GGGGACCACTTTGTACAAGAAAGCTGGGTATCAATTAAGTTTGTGCCC, 3’-GGGGACCACTTTGTACAAGAAAGCTGGGTATCAATTAAGTTTGTGCCC). The synaptophysin-pTagRFP PCR product was then subcloned into a middle entry vector, and then ultimately subcloned under a 10x element UAS promoter to generate UAS-Synaptophysin-pTagRFP.

#### Single-neuron labeling

To label V2a neurons, alx:Gal4 was co-injected with either UAS:mcd8GFP or UAS:ptagRFP plasmids into the one-to four-cell-stage in wildtype or in Tg[GlyT2:lRl-Gal4;UAS:GFP] x Tg[Dbx:cre] double transgenic embryos using a microinjector (Narishige model IM300). To label V0d neurons, Tg[GlyT2:lRl-Gal4;UAS:GFP] x Tg[Dbx:cre] zygotes were injected with either UAS:mcd8GFP or UAS:ptagRFP plasmids. Alternatively, GlyT2:Gal4 was co-injected with either UAS:mcd8GFP or UAS:ptagRFP plasmids in wildtype zygotes. Both methods labeled morphologically indistinguishable types of neuron. For labeling V0d neurons and their putative synaptic output in relation to motor neurons GlyT2:Gal4, UAS:mCerulean and UAS:Synaptophysin-pTagRFP were co-injected into zygotes from an enhancer trap line which labels axial motor neurons with GFP (Balciunas et al., 2004).

#### Electrophysiology

Zebrafish larvae were first immobilized using extracellular solution containing *α*-bungarotoxin (0.1% w/v; composition in mmol/L: 134 NaCl, 2.9 KCl, 2.1 MgCl_2_, 10 HEPES, 10 glucose, 2.1 CaCl_2_, adjusted to pH 7.8 with NaOH) and then transferred to a Sylgard-lined glass-bottom dish containing toxin-free extracellular solution. Larvae were pinned down through the notochord using custom sharpened tungsten pins then the skin was carefully removed using fine forceps. For paired recordings, muscle segments overlying the spinal cord were carefully dissected away using tungsten pins and fine forceps. Putative post-synaptic targets were sampled within neighboring spinal segments up to four segments away. Following the dissection, the preparation was moved to the recording apparatus (AxioExaminer, Zeiss) equipped with a 40x/1.0 NA water immersion objective and three motorized micromanipulators (Scientifica).

For whole-cell recordings, standard wall glass capillaries were pulled to make recording pipettes with resistances between 5-15 MΩ, which were then back-filled with patch solution (composition in mmol/l: 130 K-gluconate, 2 MgCl_2_, 0.2 EGTA, 10 HEPES, 4 Na_2_ATP, adjusted to pH 7.3 with KOH) containing either Alexa Fluor 488 or 568 hydrazide (final concentration 50 μmol/l) to visualize cell morphology at the end of experiments using a cooled CCD camera (Rolera-XR, Q-Imaging). Images were captured using Qcapture Suite imaging software (Q-Imaging). Electrophysiological recordings were acquired using a Multiclamp 700B amplifier, a Digidata series 1322A digitizer, and pClamp software (Molecular Devices). Standard corrections for bridge balance and electrode capacitance were applied in current-clamp mode. Electrical signals from spinal cells were filtered at 30 kHz and digitized at 63 kHz at a gain of 10 (feedback resistor, 500 MΩ). Connectivity was assessed by delivering 5ms step pulses at a low frequency (2 Hz) to elicit a single spike in the presynaptic cell while assessing postsynaptic responses in current clamp mode and voltage clamp mode (holding potential, –65mV).

To simultaneously monitor ‘fictive’ motor activity, in some experiments a third electrode was used for peripheral motor nerve recordings (n = 71 of 191). Pipettes were fashioned from the same ones used for whole-cell recordings; however, the tip was cut to make a 20–50 μm opening and fire polished to bend the electrode to accommodate for the approach angle. To stimulate ‘fictive’ swimming, a tungsten concentric bipolar electrode was lowered onto the tail skin using a manual micromanipulator (YOU-3; Narishige) and a brief electrical stimulus (5-20 V; 0.2 – 0.4 ms) was delivered via an isolated stimulator (DS2A-Mk.II; Digitimer). Extracellular signals from the peripheral motor nerves were amplified using a differential AC amplifier (model 1700; A-M Systems) at a gain of 1000 and digitized using Digidata 1322A with low-and high-frequency cutoffs set at 300 and 5000 Hz, respectively. The recruitment pattern during fictive swimming for V2a and V0d neurons was assessed in current clamp mode while simultaneously recording peripheral signals. For synaptic event examination, the same intracellular solution as for current clamp recordings was used in voltage clamp mode (holding potential, –65mV). No series compensation was used and the average series resistance for the cells included in the analysis was calculated as 42.4 ± 18.2 MΩ (n = 31).

For paired soma-axon recordings, we screened for fish with sparse V2a neuron GFP expression using DNA microinjections as described above. Once a stable whole-cell recording was established at the soma, fluorescence with a high-attenuation filter was used to target the axon. A loose-patch seal was achieved by applying light suction (<15 mmHg) onto the axon using glass electrodes (4-8 MΩ resistance) filled with extracellular solution. Axonal signals were recorded in current-clamp mode. For re-patch recordings, whole-cell recordings were first achieved between a V2a neuron and a motor neuron. Once the nature of the connection was tested in current-clamp mode, the morphology of the motor neuron was then captured. Afterwards, the electrode from the first recorded motor neuron was retracted and a second motor neuron was then targeted with a new electrode within the same segment while maintaining the V2a neuron recording. The same protocol was repeated to assess synaptic connectivity and confirm motor neuron morphology.

For pharmacological experiments to test the nature of connectivity, the glutamate receptor antagonist NBQX (Abcam) was dissolved in extracellular solution and delivered to the perfusate (10 μM final concentration) by a gravity-fed perfusion system. The gap junctional blocker 18-beta-glycyrrhetinic acid (Sigma-Aldrich) was first dissolved in DMSO to obtain a stock solution at 200 mM and was then used at a final concentration of 100-150 μM in extracellular solution.

#### In vivo confocal microscopy

To image transgenic lines and stochastically labeled neurons, larvae were first anesthetized in a 0.02% solution of MS-222 (Sigma Aldrich) and then embedded in low melting point agarose (1.4% in system water) in a glass bottomed dish and covered in 10% Hank’s solution with anesthetic. Fish were oriented such that the spinal cord was imaged from a lateral view. Image Z-stacks were acquired with a confocal microscope (Zeiss LSM 710) using a Zeiss 20X/1.0-NA water-immersion objective. Blue, green, and red fluorophores were excited with 458 nm, 488 nm and 543 nm laser lines respectively. Images for different wavelengths were collected sequentially to minimize signal bleed through across fluorophores. These data were collected concurrently with a DIC image of the spinal cord to locate the dorsoventral position of the neuron with respect to the top and bottom edge of the spinal cord.

#### Quantification and statistical analysis

All electrophysiological data were analyzed off-line using Igor Pro 6.2 (Wavemetrics) and organized in Microsoft Excel. The intrinsic firing pattern of V2a and V0d neurons was determined by applying 500-ms depolarizing current pulses up to a maximum of 3X rheobase. Rheobase was determined as the least amount of current to elicit spiking. Input resistance was determined by taking the average of at least three 500-ms-long hyperpolarizing pulses (–10-50 pA) within a linear range of the current–voltage relationship. Instantaneous spike frequencies were calculated from the interval of the first two spikes. For bursting neurons, burst frequency was measured by taking the interval between the peak of the first spike in each burst. Frequency values for bursting are only reported when they could be unambiguously identified by eye (typically up to 1.2X rheobase).

The conduction velocity of V2a cells was determined by measuring the distance between the somatic and axonal recording sites and dividing by time difference between the peak of the spike *dv/dt* and the peak of the axonal response. The distance between recording sites was measured using ImageJ by tracing the labeled axon in 2D from the base of the soma to the tip of the axonal electrode (mean recording distance: 389.1 ± 49.6 μm, n = 11). Three distance measurements were taken and the average distance value was used for the conduction velocity calculation. The amplitude of the axonal spike was measured from peak to trough. Fluorescent and DIC images captured at the end of each recording were used to confirm neuronal identity. Soma cross-sectional area and dorsoventral soma position normalized to the top and bottom edges of spinal cord were calculated from differential interference contrast images using ImageJ software.

To assess synaptic connectivity between cells we aligned presynaptic spike sweeps to the peak of the spike (10-200 sweeps per paired recording). Postsynaptic responses (EPSPs and EPSCs) were smoothed (5 point boxcar averaging, 10 KHz sampling rate) and the amplitude of the response was calculated by subtracting the averaged baseline value taken 2 ms prior to presynaptic spike from the peak response value. In order to get an accurate amplitude measurement of the second glutamatergic component in mixed synaptic responses, the early electrical component was subtracted out. This was achieved by taking the average from electrical responses where the glutamatergic component was visibly absent and subtracting it from all the sweeps in that trial.

Rise time was calculated from the 10-90% of the peak amplitude. Decay time for electrical components (slow and fast) was calculated by a single exponential fit between peak amplitude and baseline of the response. For glutamatergic synaptic events, the decay time was calculated by a double exponential fit between peak and 20% of peak amplitude. The glutamatergic component decay tau was taken as the first constant *τ*1 from the fit, given the potential confound of serial, non-impulse mediated interactions. Connection probability was determined by counting the number of recordings where any given synaptic was present and dividing by the total number of recordings. Synaptic failures were determined by dividing the number of responses without a fast synaptic event by the total number of spike-triggered events.

For synaptic event detection during fictive swimming, a threshold value of 6 pA was used to detect synaptic events (EPSCs) in Igor Pro 6.2 using the spontaneous activity analysis (SpAcAn) interface as described previously (Bagnall and McLean, 2014). The peripheral nerve recording was used to determine the swim frequency and then each synaptic event was assigned a frequency range using the peripheral recording as a reference. For the synaptic event detection at slow (20-30 Hz) and fast (40-50 Hz) swim frequencies, we used experiments in which the peripheral recording was only 1-2 body segments away from the recorded neuron and analyzed a minimum of 5 swim bouts and 3 swim bursts per frequency bin. Synaptic currents were analyzed from at least 5 swim bouts per fish to obtain the peak excitatory current.

The coupling coefficient between two electrically coupled neurons was calculated as the ratio of membrane potential deflections (Δ*V2*/Δ*V1*) in response to a 500-ms hyperpolarizing current pulse. Δ*V1* is the change in membrane voltage in the cell receiving the hyperpolarizing step pulse and Δ*V2* is the change in membrane voltage in the coupled cell.

To analyze firing reliability at different speeds of swimming, swim bursts were binned at 10 Hz and the likelihood of spiking (expressed as percent of total bursts) at any given speed bin was determined for each cell. The minimum swim frequency was the average from the three lowest swim frequencies at which any given cell was recruited. The spike timing of V2a neurons relative to motor neuron spiking was determined only from paired recordings from which both cells were located within the same segment. For each motor neuron spike in a given burst, the relative spike timing in reference to the motor neuron was calculated by subtracting the motor neuron time from each of the V2a spikes. These V2a spike times were binned at 0.25 ms and a histogram was constructed expressed as a percent of total spikes. The histogram was fitted to a Gaussian function and the averaged Gaussians and standard error were then plotted.

All data were tested for normality and the appropriate statistical analysis was performed using the StatPlus plug-in for Microsoft Excel. Data are reported as means plus/minus standard deviation, with corresponding tests, critical values and degrees of freedom, unless noted otherwise.

KEY RESOURCES TABLE
| REAGENT or RESOURCE | SOURCE | IDENTIFIER |
| --- | --- | --- |
| Antibodies |  |  |
| None |  |  |
| Chemicals, Peptides, and Recombinant Proteins |  |  |
| None |  |  |
| Deposited Data |  |  |
| None |  |  |
| Experimental Models: Organisms/Strains |  |  |
| Zebrafish Tg[Chx10:GFP] | Kimura et al., 2006 | N/A |
| Zebrafish Tg[Chx10:IRI:GFP] | Kimura et al., 2006 | N/A |
| Zebrafish Tg[MNET2:GFP] | Balciunas et al., 2004 | N/A |
| Zebrafish Tg[GlyT2:GFP] | McLean et al., 2007 | N/A |
| Zebrafish Tg[Dbx:Cre] | Satou et al., 2012 | N/A |
| Zebrafish Tg[GlyT2:IRI:Gal4;UAS:GFP] | Satou et al., 2013 | N/A |
| Recombinant DNA |  |  |
| UAS:mCD8GFP | Tol2kit, Kwan et al., 2007 | N/A |
| UAS:pTagRFP | Menelaou et al., 2014 | N/A |
| GlyT2:Gal4 | Satou et al., 2013 | N/A |
| UAS:mCerulean | This paper | N/A |
| UAS:Synaptophysin-pTagRFP | This paper | N/A |
| Software and Algorithms |  |  |
| None |  |  |
| Other |  |  |
| None |  |  |

